# Network analyses reveal D clade ethylene response factors as major regulators of jasmonic acid-mediated resistance to early blight disease complex in tomato

**DOI:** 10.1101/2023.10.14.562343

**Authors:** Christopher Tominello-Ramirez, Lina Muñoz Hoyos, Mhaned Oubounyt, Remco Stam

## Abstract

Resistance mechanisms to early blight disease complex (EBDC) in tomato remain obscure given its polygenic and quantitative nature. We investigated the early defense responses of Heinz 1706 tomato to EBDC using RNA-seq. We observed distinct transcriptional reprofiling upon exposure to two EBDC isolates and the PAMP chitin. Avirulent isolate CS046 (*Alternaria alternata*) elicited a vigorous defense response in the host, whilst the virulent isolate 1117-1 (*Alternaria* sect. *Porri*) showed subdued gene expression, suggesting a suppression of defense responses during compatible pathogenesis. We emphasize the specific roles of *ETHYLENE RESPONSE FACTORs* (*ERF*s) in defense against EBDC, with a particular focus on the D clade *ERFs*. Co-expression network analysis revealed the principal genes in early defense responses to EBDC are secondary metabolite biosynthesis genes, transcription factors, and hormone response genes. We constructed a gene regulatory network and predicted novel hub genes as putative global regulators of the defense response, including the D clade *ERFs, WRKY*, and *NAC* transcription factors. Our work highlights the failure of virulent EBDC pathogenesis to elicit hormone responses that suppress cell death. Additionally, we found a selective induction for specific ERFs that strongly influence the topology of the EBDC defense transcriptional network.

## Introduction

The domestication of tomatoes has eroded its genetic diversity compared with its closest wild relatives [1]. As a consequence, tomato yields are at significant risk from all forms of biotic stress, including over 200 diseases [2]. Biotic stress reduces global tomato yields by 34% under current management practices; weeds, herbivores, and microorganisms are responsible for yield losses of 6%, 10%, and 18% respectively [3]. *Alternaria* spp. are the causal agents of early blight and brown leaf spot, often cited as the most important foliar pathogens of tomato [3, 4]. Consistent with the disproportionately high representation of *Alternaria* spp. in global soils [5], phytopathogenic *Alternaria* spp. are found on every continent and in every region of major tomato production [6]. Early blight is caused by large-spored *Alternaria* sect. *Porri* [7], specifically *Alternaria linariae* (Neerg.) E.G.Simmons, commonly synonymized as *Alternaria tomatophila* E.G.Simmons or the potato pathogen *Alternaria solani* Sorauer [8]. Brown leaf spot is caused by small-spored *Alternaria* sect. *Alternaria* [7], many of which are now assigned to the *Alternaria arborescens* E.G.Simmons species complex or synonymized into *Alternaria alternata* (Fr.) Keissl. [9], a pathogen known to infect over 100 host species [7]. Brown leaf spot and early blight are a symptomatology continuum of small dark-brown lesions developing on leaves, stems, and fruits that develop characteristic concentric rings, about 5 mm in diameter for brown leaf spot and about 15 mm in diameter for early blight, culminating in complete defoliation of the plant if left unmanaged [7]. Both diseases will be referred to hereafter as early blight disease complex (EBDC) because of the significant overlap in disease severity [10] and reports that small-spored *Alternaria* spp. can also cause early blight in tomatoes [11, 12].

Due to an absence of tomato varieties sufficiently resistant to EBDC, control measures consist of cultural practices and fungicide use [13]. There are several commercially available tomato cultivars moderately resistant to EBDC such as the ‘NC EBR’ lines, but these resistant varieties do not constitute an alternative strategy to cultural practices or fungicide use; they merely extend fungicide intervals from 5 days to 10 days [14]. There are reports of full EBDC immunity in potato/wild potato hybrids [15], but no commercial cultivar has been developed, and its transferability to tomato is undemonstrated. Fungicide remains an effective preventative measure against EBDC, but despite reproducing clonally, *A. linariae* and *A. alternata* are considered medium-risk and high-risk pathogens respectively in developing fungicide resistance to several chemistries [16]. EBDC has been shown to independently evolve fungicide resistance mechanisms in the same genes in non-admixed genetic backgrounds [17]. Due to the environmental, financial, and biosecurity-related cost of intensive fungicide use, identification of host resistance mechanisms to EBDC remains an important albeit elusive target. Resistance is predicted to proceed quantitatively, polygenically, and inherited recessively due to the necrotrophic lifestyle of its causal agents [18, 19]. Accordingly, despite single genes implicated in an EBDC resistance locus in wild potato [15] and resistance to *A. alternata* leaf blotch of apple [20], only polygenic and quantitative EBDC resistance mechanisms have been identified in tomato [13].

Genetic mechanisms of defense against pathogen invasion begin when ligand-binding receptors located on the cell surface, known as pattern recognition receptors (PRRs), detect pathogen-associated molecular patterns (PAMPs) [21]. Detection of PAMPs initiates an immune signaling cascade resulting in pattern-triggered immunity (PTI), and in response, pathogens have evolved secreted proteinaceous effectors that prevent PTI, resulting in effector-triggered susceptibility (ETS) [22]. Reciprocal evolution has produced cytoplasmic receptors known as nucleotide-binding domain, leucine-rich-repeat containing receptors (NLRs) that bind effectors, blocking ETS in a process known as effector-triggered immunity (ETI) [22]. PTI and ETI are mediated by large-scale shifts in gene expression [23], secondary metabolism [24, 25], and tailored to specific pathogen lifestyles using phytohormones. Salicylic acid (SA) is the major phytohormone regulating resistance against biotrophs, pathogens that parasitize living tissue, and jasmonic acid/ethylene (JA/ET) for necrotrophs, pathogens that mobilize nutrients from dead tissue [19]. The gene-for-gene innate immunity model is predicted to invert for necrotrophic pathosystems due to the tendency for a hypersensitive response (HR) to promote defense to biotrophs and virulence in necrotrophs [19]. SA is necessary and sufficient to induce HR in some contexts [26, 27], but SA defense-signaling promotes virulence in necrotrophs even in the absence of HR [28], due to the well known reciprocal antagonism of SA- and JA-induced defense signaling [29]. Exogenous SA promotes biotroph defense at the expense of necrotroph susceptibility by antagonizing JA signaling via a myriad of mechanisms [30], and exogenous JA performs oppositely [31]. This simple paradigm predicts that SA should promote EBDC infection, and JA should suppress it [19]. Indeed, foliar application of methyl jasmonate methyl ester does reduce EBDC symptoms by 60% [32], yet unexpectedly, foliar application of SA has been shown to reduce the occurrence of lesions caused by EBDC without any intrinsic antifungal activity [33]. This benefit of priming tomato plants with SA likely arises from the induction of PATHOGENESIS-RELATED (PR) proteins [34], as *PR* transcripts have been shown to be upregulated in response to EBDC [35], and resistant lines have higher basal expression of *PR* genes [36]. Nevertheless, EBDC resistance has been linked to PR protein-induced HR, with the hypothesis that a functional difference between toxin-induced cell death and the programmed cell death of HR underlies the resistance [36]. Even JA itself can promote toxin-induced cell death from EBDC [37], but HR does not necessarily antagonize JA signaling in certain contexts [28]. SA/JA antagonism is relaxed in the presence of ET [38], highlighting a key role for ET to prioritize JA signaling. ET is an early response gene in plant immunity systems and major regulatory hub [39]. ET is a critical defense component in JA-regulated defense signaling [40, 41].

To begin disentangling the complex molecular participants of the critical early events in EBDC resistance in tomato, we sequenced transcriptomes of Heinz 1706 tomatoes [42] challenged with EBDC from virulent and avirulent isolates of EBDC and the PAMP chitin for 3 hours. The isolate CS046, a specimen of *A. alternata*, was collected *in situ* from wild tomatoes in Peru [43] and shown to be avirulent on Heinz 1706 [44]; the isolate 1117-1, a specimen of *Alternaria* sect. *Porri* was collected from tomatoes in Germany [43] and shown to be virulent on Heinz 1706 [43, 44]. Since hormones play an important role in EBDC defense, large-scale pleiotropic gene expression remodeling is expected. To identify a defense-related module of genes amongst the diverse genes regulated by the hormone flux, we constructed a weighted gene co-expression network and identified a subnetwork associated with EBDC defense. To identify the genes with the largest influence on the defense subnetwork topology, we identified hub genes with high eigencentrality and constructed a directed gene regulatory network (GRN). From these genes, we identified novel genes with roles in EBDC defense.

## Methodology

### Sample growth conditions, experimental treatment, and sample collection

Our Heinz 1706 tomatoes were grown in a phytotron-type walk-in growth chamber at 23°C with a 12-hour photoperiod. All treatments were performed on 3-week old meristem cuttings propagated from 6 week old seedlings. The fungal isolates CS046 [43] and 1117-1 [44] were grown in a Sanyo MLR-351H Versatile Environmental Test Chamber (Moriguchi, Japan) growth cabinet at 23°C, 60% relative humidity, and a 24-hour photoperiod in both the visible and ultraviolet range. Fungi were propagated on synthetic nutrient agar medium [45] in standard 100mm petri plates. Conidia were harvested from plates approximately one month after inoculation by flooding the plate with sterile millipore-filtered water. The yield of conidia was quantified using a haemocytometer, and diluted to achieve 3×10^4^ conidia×1mL^-1^ for plant infection. Crab shell chitin was prepared by freezing it in liquid nitrogen and grinding it with a mortar and pestle until a very fine powder was achieved. We spray-infected the above-ground tissues of the tomatoes with the conidia suspension until inoculum runoff was achieved, indicating droplet saturation. The same treatment method was used for 50μg chitin×1mL^-1^, and with sterile millipore-filtered water. Each treatment had 4 biological replicates. Plants were placed in a plastic tub with a lid on to ensure 100% relative humidity, as measured with an AHT20 temperature and humidity sensor (Adafruit, New York City, NY, USA). We collected treated leaves after 3 hours incubation, wrapped them in aluminum foil, flash-froze them in liquid nitrogen, and stored them at -80°C prior to RNA extraction.

### RNA extraction, library construction, and read mapping

We extracted total RNA using RNeasy Plant kits (Qiagen, Venlo, Netherlands). The mRNA library for 3’ sequencing was generated using the QuantSeq 3’mRNA-Seq Library Prep Kit (Lexogen, Vienna, Austria) and then sequenced using an HiSeq2500 (Illumina, San Diego, CA, USA) with Rapid SBS v2 chemistry to generate 100bp single-end reads. For all RNA extraction, quality assessment methods, mRNA library generation, and sequencing utilized the manufacturer’s recommended methodologies unchanged. We used ‘Trimmomatic’ [46] for initial quality filtering and adapter trimming of the raw sequencing reads, aligned trimmed RNA-seq reads to the International Tomato Annotation Group (ITAG) genome version 4 [47] using ‘HISAT2’ [48], and quantified aligned sequencing reads using ‘featureCounts’ [49]. All software used the default settings except the following Trimmomatic settings: LEADING:3, TRAILING:3, SLIDINGWINDOW:4:15, and MINLEN:40.

### Differential gene expression analysis

We used the R package ‘DESeq2’ ver. 1.38.3 [50] to perform the differential gene expression analysis. The transcript counts matrix was pre-filtered for genes with less than 10 total normalized read counts across all samples prior to differential expression analysis. Log_2_ fold change (LFC) values were shrunken using the ‘apeglm’ method [51] to reduce noise [50], and p-values were corrected using the Benjamini-Hochberg method to calculate their false discovery rates (FDR) [52]. We considered genes to be differentially expressed if their |LFC| > 1 and FDR < 0.05. All subsequent analyses utilized the normalized gene counts matrix with a regularized log_2-_transformation to reduce heteroscedasticity. The upset plot was generated using the R package ‘ComplexHeatmap’ ver. 2.14.0 [53], and the PCA was generated using DESeq2.

### GO term enrichment

We used the Cytoscape ver. 3.10.1 [54] app BiNGO ver. 3.0.5 [55] for GO term enrichment and visualization using the hypergeometric statistical test and FDR correction. GO term annotations were acquired primarily through ITAG resources. Any remaining genes without GO terms were annotated using PANNZER2 ver. 15.12.2020 [56] with default settings, and filtered for hits with a positive predictive value (PPV) above 0.4 [57].

### Analysis of canonical immunity elements

Receptor-like kinases (RLKs) were identified using ‘DeepLRR’ ver. 1.01 [58] searching for the ‘LRR_RLK’ protein type, and appended this list with RLK genes from Sakamoto *et al*. [59]. Hits were filtered for subcellular location type ‘Cell membrane’ using the ‘DeepLoc’ ver. 2.0 web service using high-throughput parameters [60], tested for transmembrane domains using the ‘deepTMHMM’ web service [61], and ligand-binding domains and kinase domains were identified using HMMER ver. 3.3.2 [62] and the ‘CD-search’ ver. 3.20 web service [63]. Receptor-like proteins (RLPs) were identified in a similar manner by using the ‘LRR_RLP’ protein type option of DeepLRR and appending this lists with RLP genes from Kang and Yeom [64], hits were filtered for the ‘Cell membrane’ subcellular location using DeepLoc, transmembrane domains identified using deepTMHMM, GPI-anchor domains identified using the ‘NetGPI’ ver. 1.1 web service [65], and ligand-binding domains classified using HMMER and CD-search. Receptor-like cytoplasmic kinases (RLCKs) were identified by filtering the list of RLCKs from Sakamoto *et al*. [59] for subcellular location ‘cytoplasm’ using DeepLoc and the ligand-binding and kinase domains analyzed using HMMER and CD-search. Nucleotide-binding domain and leucine-rich repeat proteins (NLRs) were identified by filtering HMMER hits for ‘NB-ARC’, and examining the leucine-rich repeat (LRR) domains and N-terminal domains for completeness and structure using CD-search. To identify further canonical immunity genes, HMMER was used to conduct an ‘hmmscan’ on the ITAG4.0 tomato genome [42]. Calcium-responsive proteins and respirator burst oxidase homolog (Rboh) proteins were identified by filtering HMMER hits for ‘EF-hand’ and ‘Ferric_reduct’, respectively, and using CD-search to analyze the protein domains. Mitogen-activated protein kinases (MAPKs) were referenced from the list provided by Wu *et al*. [66]. Transcription factors (TFs) were referenced by the list provided by PlantTFDB5.0 [67]. Auxin responsive genes were identified by filtering HMMER hits for ‘f-box’, ‘AUX_IAA’, ‘Auxin_resp’, ‘Auxin_inducible’, ‘GH3’, ‘Mem_trans’, and ‘Aa_trans’ to identify Transport Inhibitor Response 1/Auxin Signaling F-Box, Aux/IAA, auxin response factors, small auxin up RNAs, GH3, PIN, and AUX/LAX proteins, respectively. Abscisic acid (ABA) responsive genes were identified by filtering HMMER hits for ‘Polyketide_cyc2’, ‘Pkinase’, ‘bZIP’, ‘PP2C’, and ‘SLAC1’ to identify PYR/PYL/RCAR ABA receptors, SnRK2 ABA-activated protein kinases, ABF/AREB/ABI5 TFs, ABI1/ABI2 proteins, and SLAC1/SLAH ion channels, respectively. Cytokinin responsive genes were identified by filtering HMMER hits for ‘Response_reg’ and ‘Hpt’ to identify AHK2-4/CRE1 histidine kinase receptors and HPt histidine phosphotransfer proteins. ET responsive genes were identified by filtering HMMER hits for ‘Gamma-thionin’ to identify defensins, referencing the list provided by Liu *et al*. [68] to identify ET receptors, constitutive triple response, ET insensitive 2 (EIN2), EIN3, EIN3-like, EIN3 binding F-box, ACC synthase, and ACC oxidase proteins, and by filtering the PlantTFDB5.0 TF list for ‘ERF’ to identify ethylene response factors [67]. Gibberellic acid (GA) responsive genes were identified by filtering HMMER hits for ‘Hormone_Rec’, ‘GRAS’ and ‘f-box’ to identify gibberellin insensitive dwarf1 GA receptors, DELLA proteins, and SLY1/SNE proteins, respectively. Jasmonic acid (JA) responsive genes were identified by filtering HMMER hits for ‘JAZ’, bHLH-MYC_N’, ‘NINJA’, and ‘LisH_TPL’ to identify jasmonate ZIM-domain proteins, MYC TFs, novel interactor of JAZ proteins, and TOPLESS/TOPLESS-related genes, respectively. The canonical coronatine insensitive 1 gene was identified using NCBI resources. Salicylic acid (SA) responsive genes were identified by filtering HMMER hits for ‘BTB’, ‘DOG1’, ‘Chorismate_bind’, and ‘Patatin’ to identify non-expressor of PR genes (NPR) 1-3 proteins, isochorismate synthase 1, and enhanced disease susceptibility 1 proteins, respectively. Brassinosteroid responsive genes were identified by filtering HMMER hits for ‘bHLH-MYC_N domain’ and ‘14-3-3’ to identify brassinazole-resistant1 and 14-3-3 proteins respectively. Pathogenesis-related genes were considered true PR genes if they were i) identified as orthologous using ‘orthofinder’ software ver. 2.5.5 [69], ii) shared all conserved domains using CD-search, iii) had their best NCBI BLAST [70] hits against the ‘ref_seq’ database as the originally published PR gene [34], and were iv) within the same genetic clade as their confirmed orthologs in other taxa in a fastest minimum evolution phylogenetic tree using NCBI COBALT [71]. Genes were plotted using the R package ‘ggplot2’ ver. 3.4.2 [72].

### Weighted gene co-expression network analysis

We used the R package ‘WGCNA’ ver. 1.72-1 [73] to construct a weighted gene co-expression network. The input matrix for network generation was variance stabilized and filtered for low-expression genes by determining the threshold when the normalized counts increase exponentially from the median value as calculated using the R package ‘segmented’ (mean normalized expression < 3.501). The input matrix was similarly filtered for genes with low variance (variance < 0.027). After appending our list of high-variance and high-expression genes with the remaining differentially expressed genes (DEGs) that were filtered out, the gene expression of each sample was correlated to its experimental treatment class, a binary trait, and therefore a signed network was constructed using Pearson’s correlations as recommended by the author of WGCNA. All other network parameters were set to default values except the following parameters: ‘deepSplit’ set to 0 to suppress module splitting, minimum module size set to 30, and ‘mergeCutHeight’ set to 0.37 to prevent insignificantly-correlated modules of the minimum size from overcoming the module merger threshold. The network was then filtered for low edge weight (edge weight< 0.154) using the R package ‘segmented’ before visualization with Cytoscape. Hub genes were identified by calculating eigenvector centrality of individual co-expression modules using the R package ‘igraph’ ver. 1.5.0 [74]. Heatmaps were generated using the R package ‘ComplexHeatmap’.

### Gene regulatory network

After obtaining a comprehensive list of tomato TFs from PlantTFDB5.0 [67], we used the R package ‘Genie3’ ver. 1.20.0 [75] to infer a directed GRN of the entire input matrix of genes filtered for low-variance and low-expression. The resulting GRN was visualized using putative regulatory links filtered out based on low model fit and visualized using Cytoscape. GRN hub genes were identified by calculating eigenvector centrality of the entire GRN using the R package ‘igraph’. Heatmaps were generated using the R package ‘ComplexHeatmap’.

## Results

### Early responses to chitin and conidiospores of CS046 and 1117-1 each induce distinct transcriptome responses

To investigate the early defense responses to EBDC, we generated an RNA-seq dataset from Heinz 1706 tomato plants treated with an avirulent isolate CS046 (*A. alternata*), a virulent isolate 1117-1 (*Alternaria* sect. *Porri*), and chitin for 3 hours. Our yield of sequencing reads was 7.6 million reads per sample with a mapping rate of 86%, and identified 910 DEGs across all treatments. Principal component analysis shows discrete and distinct responses for all treatments, with sample replicates segregating together by gene expression patterns (**fig. 1A**). An upset plot shows that the treatment with the strongest response is CS046-treated tomatoes with 397 unique DEGs, and the weakest response is the 1117-1 treatment with 106 unique DEGs (**fig. 1B**). 83 DEGs (9%) were shared between all treatments; in contrast, 641 DEGs (72%) were unique to the specific treatments (**fig. 1B**).

**Figure 1.**
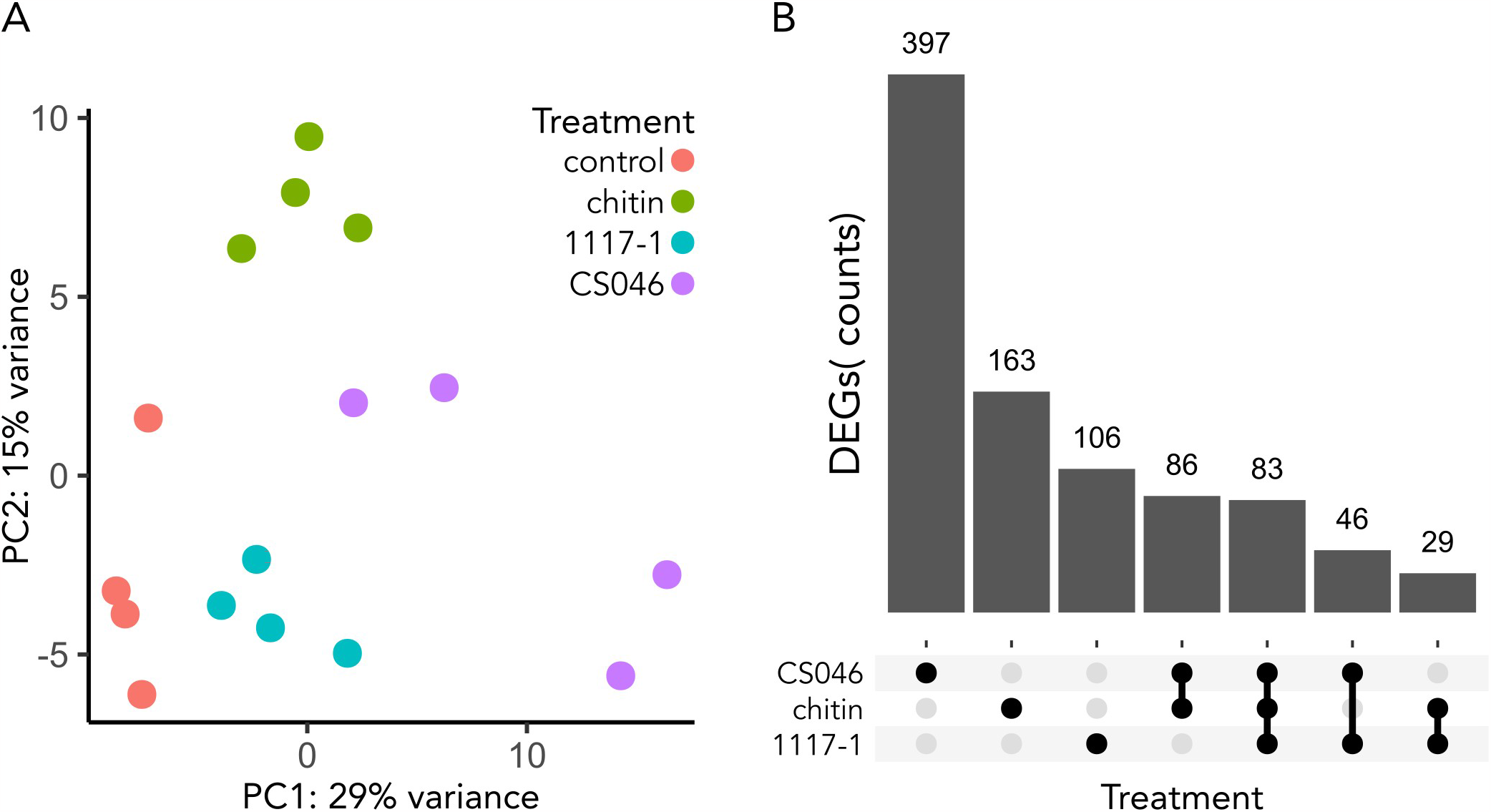
All treatments induce distinct, mostly unique transcriptome responses. **(A)** Principal component analysis plot to visualize the distribution and clustering of samples based on their gene expression. Dots represent an individual sample replicate. Dot color represents treatment class. The axes represent the variance explained by the first two principal components. **(B)** Upset plot showing the intersections and unique sets of differentially expressed genes (DEGs) amongst the treatments. The x-axis shows the categories of the comparison sets, and the y-axis represents the number of DEGs within a comparison set. The horizontal bars represent the individual treatments, the black dots indicate which treatments are compared, and the vertical lines represent the intersection of treatments for a given comparison. The numbers above the vertical bars indicate the number of DEGs in a comparison set.

### Gene expression for the CS046 treatment is more enriched for defense-related gene ontologies than for the 1117-1 treatment

To investigate the biological activities of the transcriptional responses to the treatments, we performed GO term enrichment analysis on the sets of DEGs of each individual treatment. The transcription response to chitin, 1117-1, and CS046 treatment yielded 112, 135, and 175 total enriched GO terms respectively (**supp. table 1**), indicating stronger coordination of the early defense response to the avirulent EBDC isolate. The two EBDC isolates elicit similar types of defense responses, although conspicuously missing from the 1117-1 treatment is cell wall remodeling (**table 1**). The most notable difference between the fungal treatments and the chitin treatment is the expanded enrichment of GO terms for secondary metabolite biosynthesis (**table 1**). All treatments share a common response in biological processes for ROS activities, secondary metabolite biosynthesis, stress responses, and hormone responses; molecular functions for catalytic activity, transferase activity, and molecule binding; and cellular compartments for extracellular region, plasma membrane, and the chloroplast (**table 1**). Considering the greater gene induction in the CS046 treatment (**fig. 1**) and the observation that TF activity is an enriched GO term, CS046 has a greater induction in the number of genes for each of these gene categories, despite having similar gene types in the 1117-1 treatment.

### Canonical plant immunity components suggests the importance of phytohormone signaling in early response to EBDC

To interpret the immune signaling events during EBDC defense, we investigated the activation of canonical PTI/ETI pathways [76]; calcium signaling, hormone responses, MAP kinases, NLRs, PRs, RLPs, RLKs, RLCKs, and TFs. We found 438 genes with significant differential expression in these categories, and 136 DEGs which also satisfy the log_2_ fold change threshold (**supp. table 2**). *MAPKs*, NLRs, *PRs*, RLCKs, RLKs, and RLPs only account for 32 DEGs, while the remaining 104 DEGs are split between calcium signaling, hormone responses, and TFs, indicating the importance of these three gene categories (**fig. 2**). Tomatoes treated with CS046 have the strongest immune response with 73 DEGs identified as canonical immune system components. Tomatoes treated with chitin and 1117-1 yielded 41 and 28 DEGs respectively, consistent with the global transcription responses of the individual treatments (**fig. 1**). Notably, some of the most strongly induced genes from the CS046 treatment are five *ETHYLENE RESPONSE FACTORs* (*ERFs*) from the D clade [77], which have not been previously identified with a role in EBDC defense.

**Figure 2.**
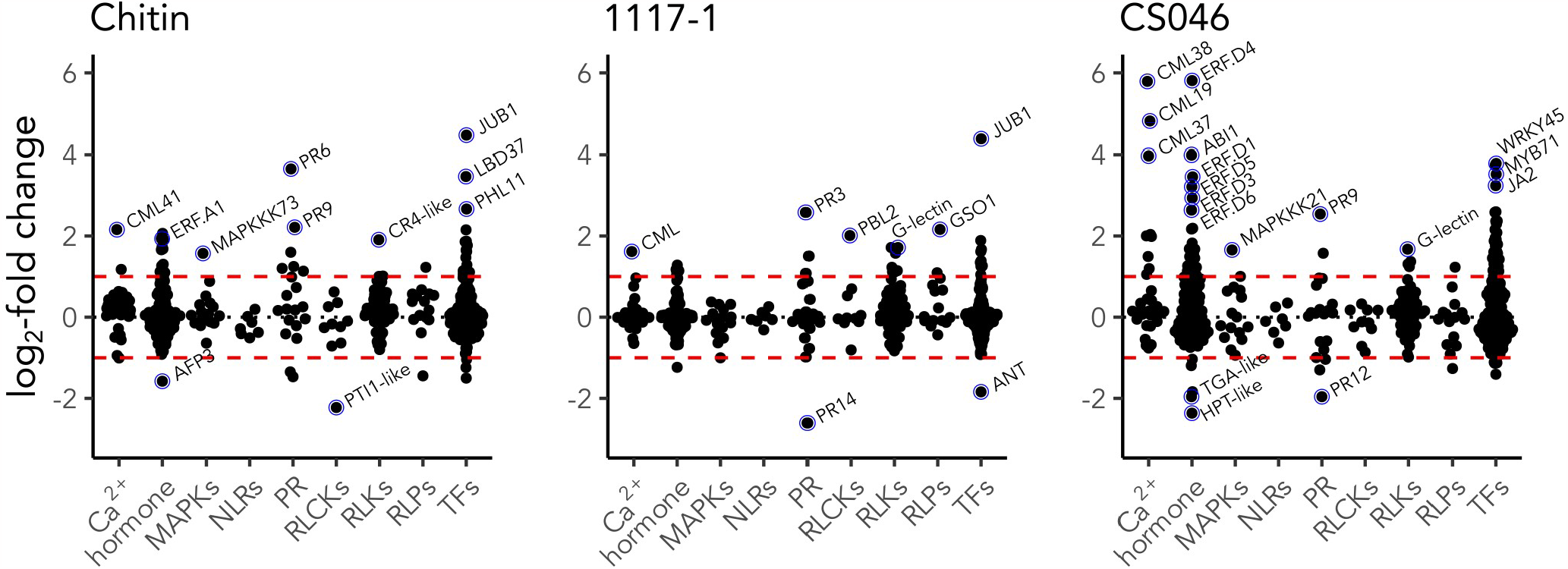
The early defense response to EBDC involves gene transcription, Ca^2+^ signaling, and hormone signaling. Expression patterns of tomato genes with significant differential expression (FDR < 0.05) in canonical immune signaling pathways after treatment with chitin or spores of either a virulent (1117-1) or avirulent (CS046) isolate of EBDC. The black dotted line is set to the up/down-regulation inflection point (LFC = 0); the red dashed line is set at the LFC threshold to be considered a DEG (LFC = |1|); blue circles and labels annotate notable genes with high differential expression.

### Defense-related co-expression network characterized by hormone signaling, cellular stress responses, and secondary metabolite biosynthesis

To identify key regulatory genes of EBDC resistance in tomato, we investigated whether the distinct transcriptional modules can be associated with specific treatments, with particular focus on a transcriptional module of defense-related genes associated with CS046 treatment. To filter out genes generally not involved in the defense responses and gain more insight into the EBDC defense response, we performed a weighted gene correlation network analysis to identify the co-expression networks underlying the CS046 defense response. 4644 genes passed our thresholds for mean expression and variance (**supp. table 3**). After filtering for edge weight, we yielded a gene co-expression network with 3430 genes placed in 8 co-expression modules, and over 200,000 co-expression relationships. The co-expression modules have distinct GO term enrichment, indicating discrete biological activities associated with specific co-expression modules (**supp. table 1**). Upon visualization the co-expression network, the boundaries between the co-expression modules are well-defined (**fig. 3A**). There are 5 significantly correlated co-expression modules with Pearson’s correlations (*r*) higher than 0.65 (*p* < 0.01) associated with each experimental treatment class; the brown module (*r*=0.93) for the mock treatment, the turquoise module (*r*=0.84) for chitin treatment, the red (*r*=0.70) and blue (*r*=0.70) modules for the CS046 treatment, and the yellow module (*r*=0.67) for the 1117-1 treatment (**fig. 3B**). Among the 2 co-expression modules that are significantly correlated to the CS046 treatment, the red module can be assigned to photosynthesis-related activities, the blue module with defense responses (**supp. table 1**). To further confirm the assignment of the blue module to the defense response, we performed a Fisher’s exact test to evaluate the enrichment of genes within the co-expression modules for genes regulated by MYC2, a master regulator of necrotrophic defense responses in tomato [78]. Genes from the blue module have 67% higher odds of being MYC2-regulated (list from Du 2017), and is the only co-expression module with statistically significant enrichment (FDR < 0.01). The blue module also has the highest average edge weight for any module in the network, with 0.24, 0.17, and 0.18 for the blue, yellow, and turquoise modules respectively.

**Figure 3.**
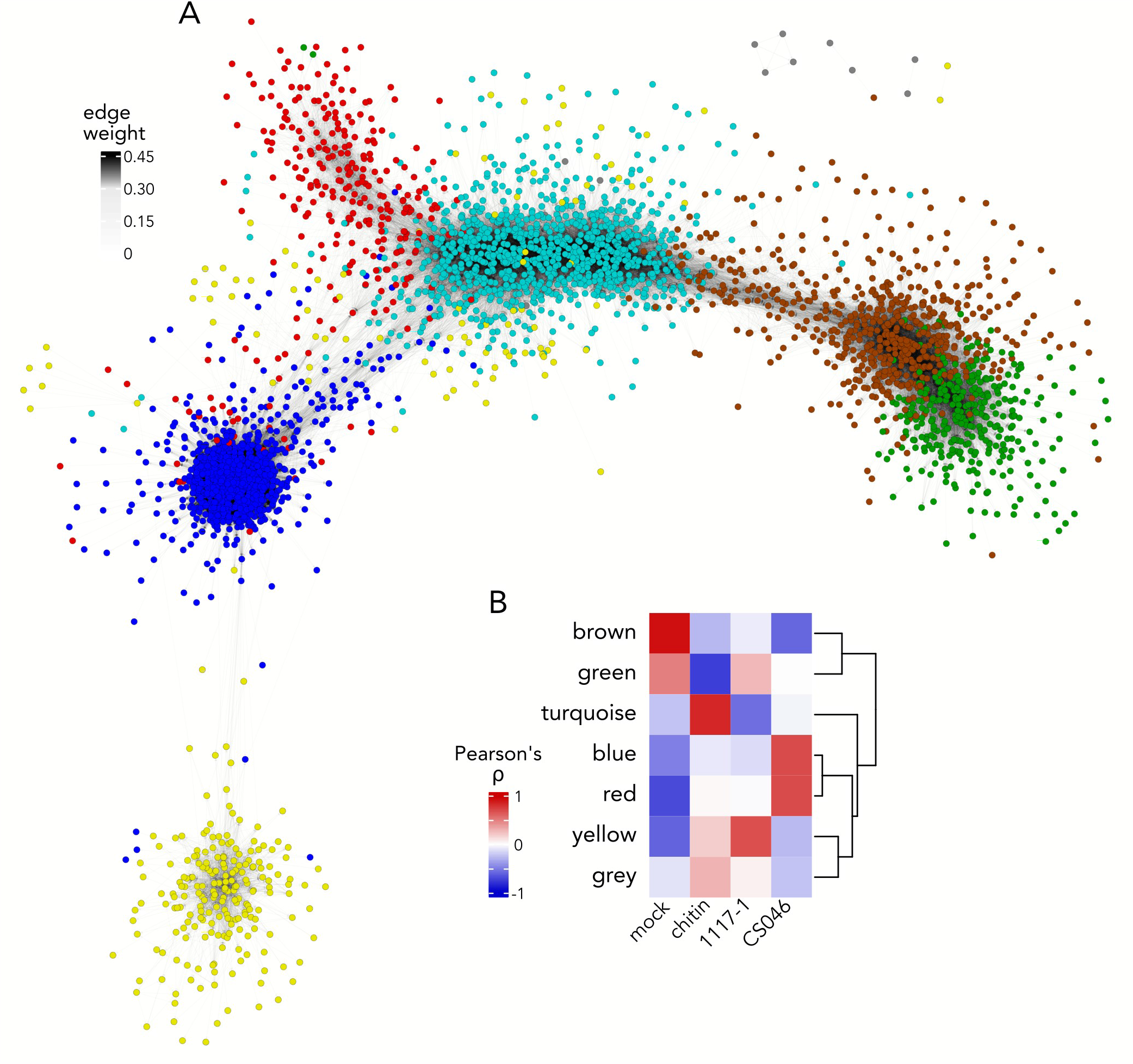
WGCNA yields a discrete co-expression module associated with EBDC defense. **(A)** Visualization of the weighted gene co-expression network analysis (WGCNA). Nodes represent genes colored according to co-expression module membership. Edges indicate a measurable co-expression relationship with weaker connections having increasingly transparent lines. **(B)** Heatmap showing module-trait relationships. Rows represent co-expression modules, and columns represent the experimental treatment. Heatmap cells are colored according to the Pearson’s correlation coefficient r between a co-expression module eigengene and an experimental treatment class. Row dendrogram illustrates hierarchical clustering results for similar co-expression modules.

### D clade ERFs are strongly upregulated exclusively in CS046 treatment and are hub genes for the defense-related co-expression module

To identify the genes with the strongest influence on the topology of the defense-related co-expression network, we calculated network eigenvetor centrality for each gene in a given co-expression module. These genes are central to the directed biological specificity of their local co-expression modules [79]. We found 20, 53, and 94 genes with high eigenvector centrality for the yellow, turquoise, and blue co-expression modules respectively (**supp. table 4**). For the yellow module’s defense-related hub genes, four genes are involved in lignin biosynthesis, three secondary metabolite biosynthesis genes, and one gene each of proteases, chitinases, ABC transporters, glutathione transferases, ABA-responsive genes, RLKs, and TFs (**supp. table 4**). Of the turquoise hub genes that are defense related, 20 genes are involved in cell wall remodeling, 8 molecule transport genes, 3 hormone response genes, 2 secondary metabolite biosynthesis genes, 2 RLK genes, and one gene each for calmodulin-binding-like proteins, protease inhibitors, lignin biosynthesis, stress responses, trichome development, and TFs (**supp. table 4**). Of the blue module genes that are defense related, 14 are associated with secondary metabolite biosynthesis, 12 TFs, 7 transport genes, 6 hormone response genes, 6 glutathione S-transferases, 6 stress response genes, 5 protein stress genes, 4 calmodulin-like genes, 4 lignification genes, 3 hormone biosynthesis genes, 3 signal transduction genes, 2 glycosyltransferase genes, 2 cell wall remodeling genes, and one gene each of ABC transporters, cuticle biosynthesis, MAP kinases, programmed cell death regulators, protein synthesis, RLKs, senescence, and wounding responses (**supp. table 4**).

TFs with high eigenvector centrality are likely to be important regulators of their respective co-expression modules. 12 of the 94 hub genes for the blue module are TFs; five *ERFs, SlERF*.*H14* (Solyc05g052410), *SlERF*.*D3* (Solyc01g108240), *SlERF*.*D4* (Solyc10g050970), *SlERF*.*D5* (Solyc04g012050), and *SlERF*.*D6* (Solyc04g071770); one myeloblastosis *(MYB), SlMYB79* (Solyc05g053150); one *WRKY, SlWRKY45* (Solyc08g067360); one homeodomain-leucine zipper (HD-ZIP), *SlHOX6* (Solyc03g082550); one *NAM/ATAF1/2/CUC2 (NAC), JA2* (Solyc12g013620); one DNA binding with one finger *(Dof), SlDof2*.*1* (Solyc06g075370); one Nuclear Transcription Factor, X-Box Binding 1 *(NFX1), NFXL1* homolog (Solyc03g118420); and one basic leucine zipper *(bZIP), SlbZIP07* (Solyc01g100460) (**fig. 4**). The yellow and turquoise modules each have one TF each amongst their hub genes, *SlWRKY16* (Solyc02g032950) and *SlERF*.*H1* (Solyc06g065820) respectively (**fig. 4**). The defense roles of the D clade *ERF*s – *SlERF*.*D3-6* – are relatively uncharacterized. It has been shown that *SlERF*.*D4-6* are upregulated during ETI of *Pseudomonas syringae* pv. *tomato* (*Pst*) strain DC3000 infection [80], and *SlERF*.*D2, SlERF*.*D6*, and *SlERF*.*D7* expression is induced in response to *B. cinerea* infection of red ripe tomato fruits but not wounding [81], suggesting a role in the defense response specific to pathogen detection. None of these TFs have been implicated in EBDC defense before, and we report a role in biotic stress in tomato for *SlbZIP07, NFXL1, SlERF*.*D3* and *SlERF*.*H14* for the first time. The D clade *ERF*s reveal themselves to be the most interesting genes to evaluate mechanistically in future studies, as they are 5 of the 6 most strongly induced hormone response genes in the CS046 treatment (**fig. 2**), they are specifically induced as a transcriptional module in CS046 treatment (**supp. table 2**), and due to the novelty of their implication in EBDC defense.

**Figure 4.**
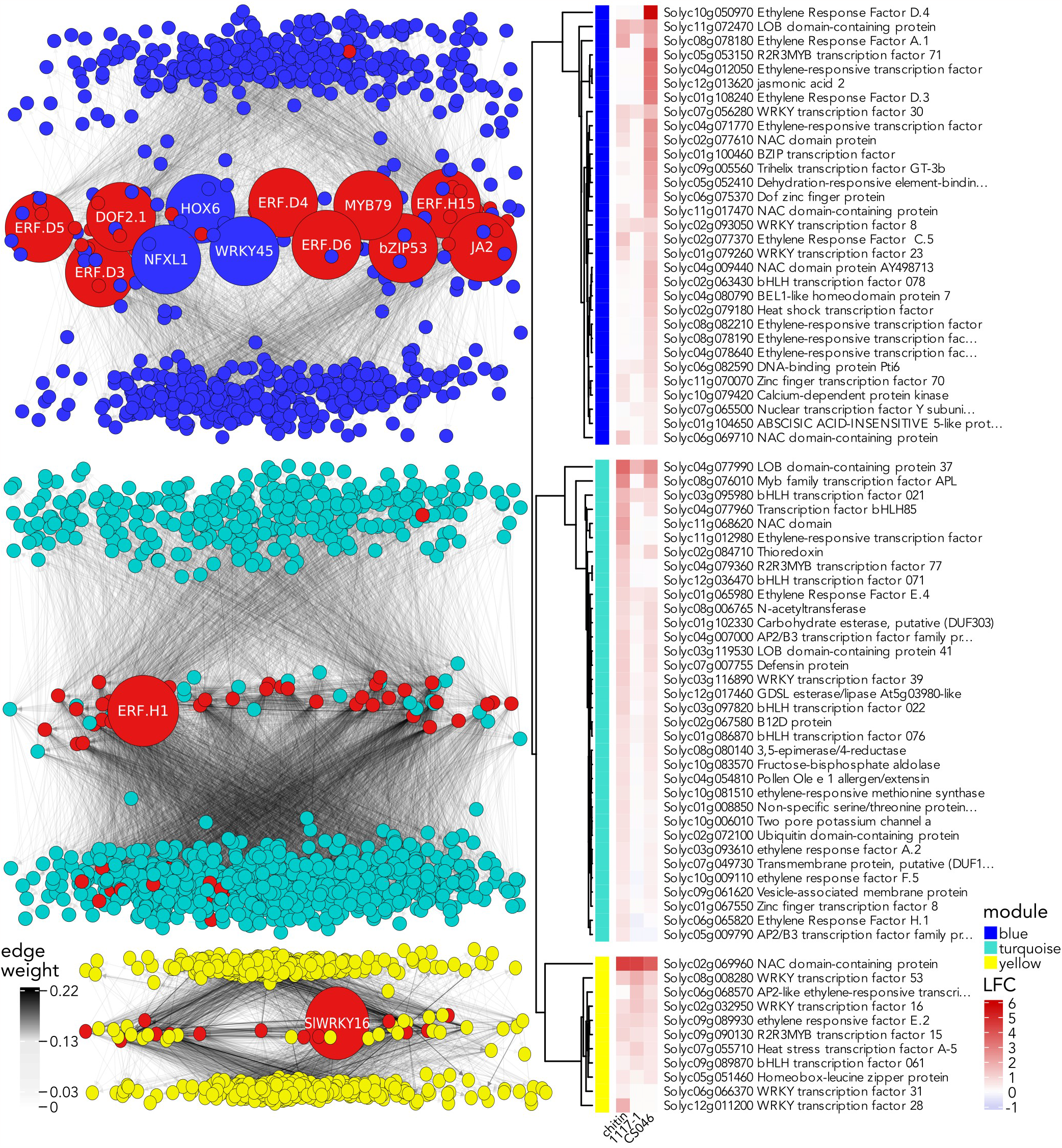
Blue module regulatory hubs show specific induction, while turquoise and yellow module regulatory hubs demonstrate generalized induction. The directional gene regulatory network (GRN) subnetworks showcase the GRN hub genes: blue for EBDC defense-associated, yellow for EBDC susceptibility-associated, and turquoise for chitin-associated co-expression modules with adjacent clustered heatmaps that are colored based on their log_2_ fold change (LFC). In the networks, nodes symbolize genes with their colors indicating module membership. Large nodes, annotated with their protein names, represent co-expression hub genes, while red nodes denote GRN hub genes.

### Gene regulatory network predicts significant functional specialization of the co-expression network hub gene transcription factors

We constructed a directed GRN using the same input matrix previously used to generate the co-expression network (**supp. table 3**) to predict potential regulatory targets of the hub gene TFs. The blue module has 12 co-expression network hub genes that are also TFs (**fig. 4**). Our GRN predicts that the six D clade ERFs regulate ∼47% of the genes within the blue module (**supp. table 5**). The GRN subnetworks regulated by each hub gene contains hundreds of genes, and therefore we used GO term enrichment to summarize their putative biological functions. All of the predicted D clade ERF gene regulatory subnetworks have enriched gene ontologies (**supp. table 1**), notably with GO terms for secondary metabolism and hormone responses being shared across all. The enriched GO terms for the *SlERF*.*D1* and *SlERF*.*D4* GRN subnetworks are similar to each other, with enriched GO terms for phenylpropanoid, coumarin, and jasmonic acid metabolism, protease and hydrolase inhibition, and responses to fungi (**supp. table 1**). Still, *SlERF*.*D1* and *SlERF*.*D4* are divergent from the other D clade *ERF*s, with uniquely enriched GO terms for brassinosteroid biosynthesis and responses to vitamin B2 in *SlERF*.*D1* and anthocyanin biosynthesis, stomata regulation, and xenobiotic transport for *SlERF*.*D4* (**supp. table 1**). The four remaining D clade *ERF*s share enriched GO terms for ET signaling, stress responses, and toxin degradation, but the more functional overlap exists between *SlERF*.*D2, SlERF*.*D5*, and *SlERF*.*D6*, which additionally share enriched GO terms for auxin responses and abiotic stress (**supp. table 1**).

Among the blue module GRN hub gene TFs, excluding the D clade *ERFs*, all have enriched gene ontologies with only GO terms for hormone responses shared between them all, largely due to *NFXL1* having GO terms for only ET signaling and stomata movement (**supp. table 1**). Excluding *NFXL1*, the remaining hub genes share enriched GO terms for toxin degradation, jasmonic acid, and phenylpropanoid biosynthesis. A particularly strong overlap exists between *JA2, SlDof2*.*1, SlbZIP07, and SlMYB79*, with additional shared enriched GO terms for ET and auxin signaling, coumarin biosynthesis, response to fungi, and sulfur homeostasis. *SlERF*.*H14* and *SlWRKY45* have additional enriched GO terms for abiotic stress in their GRN subnetworks, as well as specifically enriched GO terms for protein folding and abiotic stress for *SlERF*.*H14*, and stomatal movement and lignification for *SlWRKY45*. The turquoise module’s single hub gene that is also a TF, *SlERF*.*H1* (**fig. 4**). Our GRN predicts that it regulates ∼33% of the turquoise module (**supp. table 5**), and has enriched GO terms almost exclusively related to cell well remodeling, as well as gibberellin responses (**supp. table 1**). The yellow module’s single hub gene that is also a TF, *SlWRKY16* (**fig. 4**), is predicted in our GRN to regulate ∼23% of the yellow module (**supp. table 5**), and while its regulatory subnetwork has no significantly enriched GO terms, the most highly enriched GO terms are for biotic stress, defense responses, and toxin responses.

### Network statistics predict D clade ERFs to be global regulators of the defense-associated gene regulatory network

To identify global regulators of the GRN, we filtered the global GRN for co-expression module membership, and calculated network eigenvector centrality for each gene in a given GRN subnetwork. While these genes may not have a direct role in the biological activity of the co-expression module, but they regulate the regulators due to their basal role in the global regulatory hierarchy [79]. We calculated 40 hub genes within the blue module GRN (**supp. table 4**). The blue module is the EBDC-defense associated co-expression module, and therefore these genes likely represent specific defense responses. Many blue module GRN hub genes have well-known role in plant immunity, such as *SlERF*.*A1* (Solyc08g078180) [82], *JA2-like* (Solyc07g063410) [41], *SlERF*.*C6* (Solyc02g077370) [83], and the gene with highest eigenvector centrality of the blue module GRN, zinc finger CCCH domain-containing protein 70 (*SlC3H70*, Solyc11g070070), a gene family known to be responsive to biotic and abiotic stresses [84], but our study implicates a role in plant immunity for *SlC3H70* for the first time. 55 genes were predicted to be hub genes for the turquoise module GRN (**supp. table 4**). The turquoise module is the chitin-associated co-expression module, and therefore these genes contribute to generalized defense responses. Notable turquoise module GRN hub genes with known defense roles include three *ERF*s, *SlERF*.*H1* [85], *SlERF*.*F5* (Solyc10g009110) [86], *SlERF014* (Solyc11g012980) [87], *SlWRKY39* (Solyc03g116890) [88], defensin *SlDEF9* (Solyc07g007755) [89], and zinc-finger protein *SlZF-62* (Solyc06g075780), whose homolog in *A. thaliana* is upregulated in response to chitin [90], among others. 14 genes were predicted to be hub genes for the yellow module GRN (**supp. table 4**). The yellow module is the EBDC-susceptibility co-expression module, and therefore these genes may represent either ineffective defense strategies, or potential susceptibility factors. The gene with the highest eigenvector centrality in the yellow module GRN is *SlWRKY16* (**fig. 4**), which is predicted in our GRN to regulate every other co-expression network hub gene (**supp. table 4**). Notable defense related yellow module GRN hub genes are *SlWRKY28* [91], *SlWRKY31* [92], and *SlWRKY53* [93].

## Discussion

We investigated the early defense responses of Heinz 1706 tomato plants to EBDC using an RNA-seq dataset of tomatoes treated for 3 hours with an avirulent EBDC isolate CS046, a virulent EBDC isolate 1117-1, the PAMP chitin, and yielded disparate transcriptional profiles for each treatment. Of the 910 DEGs, 72% of them are unique to their treatments. Nearly half of all DEGs are induced exclusively in the CS046 treatment. We further characterized the early defense response using GO enrichment and network analyses. All treatments displayed GO terms enriched for classic immune responses, such as phenylpropanoid biosynthesis, ROS activities, response to toxin, and hormone responses, but only CS046 treatment induced genes annotated with gene ontologies for JA signaling, and had the strongest induction of ET-associated genes in both the strength of the induction and the diversity of the induced genes. 1117-1 treatment induced ET biosynthesis but not signaling, chitin treatment induced ET signaling but little biosynthesis, CS046 treatment induced both s. This observation potentially underscores the difference between virulent and avirulent EBDC pathogenesis; 1117-1 either abolishes ET/JA signaling, or delays it beyond 3 hours post infection (hpi). It is not sufficient to conclude that 1117-1 treatment simply fails to induce ET/JA signaling because ET biosynthesis is not impaired.

ET/JA signaling is crucial for the defense response to necrotrophic pathogens, including EBDC [19, 32]. The weak induction of canonical plant immunity components suggests compromised host defense signaling in the 1117-1 treatment. We investigated the expression of genes associated with canonical PTI and ETI signaling pathways and observed induction in almost all pathways investigated. Pathogenic effectors are often the means by which host defense signaling components can be attenuated [22], but our data does not suggest that effective EBDC defense involves induction of NLR gene transcripts to counteract an effector. If 1117-1 suppresses plant immunity with a canonical pathogenic effector, CS046 either does not possess it, or a canonical R gene is not required as an effective countermeasure against it. Hormone-responsive genes, calcium signaling, and TFs showed the greatest amplitude of induction and diversity in response among the defense gene categories we investigated. Specifically, *ERFs* seem to have consistently high induction in CS046 and chitin treated tomatoes. *SlERF*.*C1* is the only *ERF* that is a DEG in the 1117-1 treatment. It has nearly equal log_2_ fold change for all experimental treatments, and therefore has little explanatory power for the weak defense response of the 1117-1 treatment. The strong induction of *ERFs* in the CS046 treatment as a host defense strategy is consistent with our understanding of necrotrophic defense signaling [41], but the specificity of the response to the D clade *ERFs* is unreported in other pathosystems.

We generated a co-expression network to identify which TFs have the greatest influence on the topology of the transcriptional landscape during EBDC defense. We yielded one co-expression module associated with the 1117-1 and chitin treatments, and two modules associated with the CS046 treatment, the red and blue modules. The blue module is primarily associated with secondary metabolite biosynthesis and hormone signaling, whereas the red module contains photosynthesis-related genes. The red module may also represent an effective defense response, but the effects of photosynthesis modulation likely have a stochastic and pleiotropic effect on plant immunity. We infer that the blue module represents the core successful EBDC defense response. The chitin-associated turquoise module is enriched for GO terms for cell wall remodeling, and some GA signaling and mitosis. This is consistent with the effect of chitin treatment on rice, which was found to mediate resistance to a fungal pathogen by inducing compositional changes in the cell wall [94]. The yellow module is largely comprised of generic stress responses, but without ET/JA signaling, and less induction of defense-related genes in general. Assignment of the blue, turquoise, and yellow modules to defense-associated co-expression subnetworks is consistent with previous findings in *A. thaliana* that found core fungal necrotroph response genes to be RLKs, cell wall remodeling, *WRKY, NAC*, and *MYB* TFs, glycosyltransferases, calcium-signaling proteins, hormone response genes, glutathione transferases, nutrient scavenging, HR suppression, ABC1 proteins, lipid transfer proteins, heat shock factors, and thioredoxins [95], all of which are major components of these three co-expression modules.

Eigenvector centrality is a favored network statistic to predict transcription network hubs, specifically in the medical field, due to its inherent ability to detect network nodes with the largest influence on network topology, and because it is particularly adept at identifying biologically meaningful genes that other network centrality statistics fail to detect [96]. The yellow module yielded one TF as a hub gene in its co-expression network, *SlWRKY16*. It has been shown be a susceptibility factor to nematode parasitism [92], but also to increase lignification and successfully induce resistance to field dodder [97], and is upregulated during ETI of *Pst* DC3000 [80]. It is expressed in all 3 treatments however, and therefore represents either an insufficient defense mechanism, or its expression is abolished after 3 hpi by virulent EBDC so that it can no longer contribute to the defense response. The turquoise module also yielded just one TF as a hub gene, *SlERF*.*H1*, previously implicated in defense to the fungal pathogen *Rhizopus nigricans* through the expression of *PR5*, phenylalanine ammonia-lyase (PAL), and chitinase genes [85]. The turquoise module contains 2 of the 5 chitinase genes in the co-expression network, and 2 of the 3 PAL genes. PAL genes catalyze the biosynthesis of cinnamic acid and are involved in at least three different defense-related biosynthesis pathways, suberin, SA, and phenylpropanoids. Cinnamic acid and its hydroxylated derivatives display direct antifungal activity and foliar application 24 hpi reduces the severity of EBDC symptoms in tomato [98], but the genes associated with biosynthesis of cinnamic acid and its derivatives are not DEGs in any treatments, and downstream reactions that consume them are DEGs in all treatments. This suggests that at 3 hpi, these biosynthesis pathways are terminating at suberin, cuticle, and lignin biosynthesis, and accumulation of cinnamic acid derivatives to mediate EBDC defense is not justified in our analysis.

The blue module yielded 12 TF hub genes, the majority of them or their homologs have roles during of *Pst* DC3000 pathogenesis [80]. *SlbZIP07* is downregulated in response to SA, JA, and ACC [99]; *SlMYB79* is strongly induced during *Cladosporium fulvum* defense in *Cf*-12 tomatoes [100]; *SlWRKY45* is known to be a susceptibility factor for nematode parasitism [101] but participates in ET/JA signaling during fungal ergosterol/squalene perception [102]; *JA2* induces signaling to close stomata during pathogen defense [103]; *SlDof2*.*1* enhances JA-induced leaf senescence in *A. thaliana [104];* and *SlHOX6* is upregulated during ETI of *Pst* DC3000 *[80]*. The gene with the highest eigenvector centrality in the blue module GRN is *SlC3H70*, a gene family known to be responsive to biotic and abiotic stresses [84]. Many blue module GRN hub genes have well-known roles in plant immunity; the blue module GRN hub genes with the second- and third-highest eigenvector centrality are *SlERF*.*A1* and *JA2-like*, both of which directly contribute to defense to *B. cinerea* in tomatoes [41, 82], and other defense-related *ERFs*, for example, *SlERF*.*C6* [83]. Neither of these genes were identified as hub genes in the co-expression network, but 9 of the 12 genes in the co-expression network hub genes are also GRN hub genes; *SlbZIP07, SlMYB79, SlDof2*.*1, JA2, SlERF*.*H14*, and the four D clade *ERFs SlERF*.*D3-6*. Identification of these genes as local regulators of the co-expression network and global regulators of the GRN for the EBDC defense-associated transcription module highlights the potential for these genes to reveal molecular mechanisms of EBDC defense in downstream studies. The gene with the highest eigenvector centrality in the turquoise module GRN is basic helix-loop-helix 22 (*SlbHLH02*2, Solyc03g097820), one of seven ET-responsive bHLH TFs in the tomato genome [105] implicated in abiotic stress in previous studies [106]. The most highly upregulated turquoise module GRN hub gene *ERF* is *SlERF014*, a gene with that occupies the H clade of ERFs in the Pirrello *et al*. 2012 nomenclature scheme [77, 107], is known as a susceptibility factor to *B. cinerea [87]*, and is notably uninduced in EBDC-treated samples, again implicating a major role for ET triggering successful defense against EBDC. The gene with the highest eigenvector centrality in the yellow module GRN is *SlWRKY16*. In our dataset, its strongest regulatory target is a chitinase and regulates many RLKs, and several lignification genes (**supp. table 5**). *SlWRKY31* is also a yellow module GRN hub gene, and it was also found to suppress nematode defenses by Kumar *et al. [92]*. In our dataset *SlWRKY16* is expressed in all treatments and *SlWRKY31* has very low expression in general, giving neither of them explanatory power to produce susceptibility to 1117-1 and defense to CS046. A homolog of *WRINKLED3* (*WRI3*, Solyc06g068570) is a yellow module GRN hub gene, an essential gene in *A. thaliana* for cuticular maintenance by inducing cutin biosynthesis in cauline leaves [108]. The cuticle has a complex role in plant immunity [109], however it has been shown that increased cuticular lipid biosynthesis is a susceptibility factor in the *Alternaria brassicicola/A. thaliana* pathosystem [110]. Another possible yellow module GRN hub gene susceptibility factor is a homolog of *JUNGBRUNNEN1* (JUB1, Solyc02g069960), a gene known to promote leaf senescence in tomato, increasing susceptiblity to necrotrophic fungi [111]. This gene is highly expressed in all treatments compared with the mock, yet other NAC TFs that promote leaf senescence are only active in the presence of SA [112]. Due to SA/JA antagonism, senescence may be promoted in 1117-1 treatment but not in the CS046 treatment due to the indication that JA concentrations are elevated in CS046 due to the high expression of JA biosynthesis genes in the CS046 treatment. Furthermore, the most upregulated gene in the CS046 treatment is *JASMONATE-INDUCED OXYGENASE 3* (*JOX3*, Solyc10g076670), a gene that catabolizes excessive JA. *JOX3* is an indication of high levels of JA and is completely unregulated in all other treatments [113]. It is tempting to speculate that this is a major driver of 1117-1 virulence; chitin is sufficient to induce promoters of leaf senescence but it is not sufficient to induce substantial JA biosynthesis. Abolition of JA/ET pathways in the presence of PAMPs may be sufficient to derepress leaf senescence and therefore promote pathogen virulence. Previous studies have shown that an *A. solani* isolate of EBDC secretes a proteinaceous effector that increases virulence in tomato, and induces expression of senescence genes in *Nicotiana benthamiana*, primarily *SEN4, SAG12*, and *DHAR1* [114]. Orthologs of these genes are not DEGs.

Plants rapidly biosynthesize JA and ET in response to necrotrophic pathogens and herbivory, and the mechanism by which these hormones accumulate is incrementally achieving clarity. Tomato responds to herbivory with a rapid Ca^2+^ burst that activates SlCaM2 (Solyc10g081170) to bind SlERF16 (Solyc12g009240), afterwards inducing expression of itself and JA biosynthesis genes [115], increasing JA and ET by 5 and 10 fold respectively within 15 minutes [40]. *SlCaM2* has 100% sequence identity with *AtCaM7*, a calmodulin gene in *A. thaliana* that likewise induces expression of JA biosynthesis genes by derepressing the JAV1-JAZ8-WRKY51 complex [116]. Downstream JA signaling events are largely controlled by the master regulator of JA responses MYC2 [117]. In *A. thaliana*, MYC2-targeted gene activation is constitutively repressed by JAZ proteins, and derepressed in the presence of JA [117]. MYC2 also binds directly to the promoters of JA biosynthesis genes and induces their expression [118], amplifying and diversifying JA responses. It should be noted that the molecular events controlled by MYC2 in tomato are somewhat disanalogous to its homolog in *A. thaliana;* MYC2 in tomato and *A. thaliana* induces defense responses to wounding [119] but suppresses pathogen-responsive genes in *A. thaliana [120]*, as demonstrated by the increased resistance to *B. cinerea* in *myc2* [121]. In contrast, tomato MYC2 induces pathogen-responsive genes as well; overexpression lines of its regulatory targets increase resistance to *B. cinerea [41]*. In our dataset, despite low expression of JA biosynthesis genes, we find *SlCaM2/3/5* in the turquoise module with relatively high gene expression (FDR < 0.05, LFC < 1) in both the chitin and CS046 treatments. We do not however observe differential expression of *ERF16*, which should be induced by itself and JA in a feedback loop [40, 115]. These results imply that although chitin may be sufficient to initiate Ca^2+^-induced JA biosynthesis, it is either not sufficient to induce substantial JA biosynthesis, or induction of JA biosynthesis genes had ceased before 3 hpi. These results also highlight a longstanding gap in our knowledge of JA/ET signaling in defense responses – the eminent specificity of the responses. A model of the molecular events is outlined in **Figure 5**. We infer that D clade ERFs respond to EBDC specifically and similarly to SlERF16 in herbivore defenses to substantially induce JA biosynthesis genes. *ERF16* is directly induced by MYC2, and likewise so are the four D clade *ERFs* that are both co-expression network and GRN hubs, especially *SlERF*.*D3* which is both JA-responsive and MYC2-regulated *[41]*. The possibility that D clade *ERFs* respond specifically to EBDC to induce JA biosynthesis and promote resistance warrants further mechanistic studies to validate these indications.

**Figure 5.**
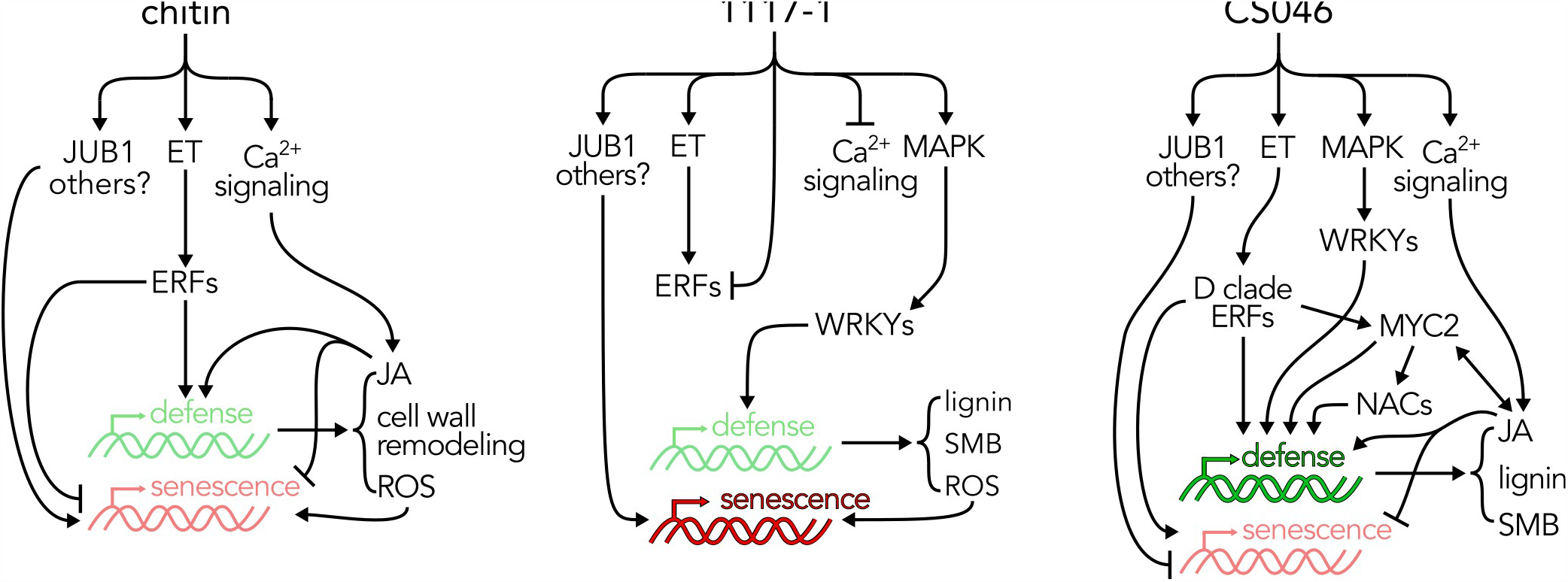
A model for D clade ERF activation of JA-activated EBDC defense responses. In tomato, chitin alone initiates immune signaling mediated by PRRs, resulting in ethylene (ET) signaling and calcium bursts. Calcium burst contributes to numerous immune processes, specifically jasmonic acid (JA) biosynthesis. ET induces the expression of *ETHYLENE RESPONSE FACTORs (ERFs)* which induce the expression of defense-related genes, including JA biosynthesis genes. Chitin also induces the expression of senescence promoting genes and the production of reactive oxygen species (ROS), which also promotes senescence and a hypersensitive response, but both JA and ERFs effectively suppress senescence. For virulent EBDC (1117-1) infection, ET biosynthesis remains intact, but ET signaling is severely suppressed. JA biosynthesis also fails to be induced, indicating the abolition of calcium bursts. *WRKY* genes are overrepresented in the GRN, which likely contributes to the 1117-1 defense response, and indicates an intact MAP kinase signaling cascade. The 1117-1 defense response has contributions from parallel pathogen detection pathways as it differs from chitin. It primarily consists of secondary metabolite biosynthesis (SMB), including lignin biosynthesis pathways. For avirulent EBDC (CS046) infection, intact ET signaling specifically induces D clade *ERFs*, contributing to substantial JA biosynthesis. D clade ERFs bind to the GCC box of MYC2-regulated genes to amplify defense responses. Though the types of genes are similar to 1117-1 treatment, the number and diversity of these genes are significantly amplified.

## Acknowledgments

This project was financially supported by the Deutsche Forschungsgemeinschaft-funded project SFB924. We gratefully acknowledge Christina Wurmser for preparing the RNA-seq libraries, Tamara Schmey for feedback and input regarding the taxonomy of the EBDC isolates used in this study and Ralph Hückelhoven and Jan Baumbach for their general support.

